# Enhanced diffusion by reversible binding to active polymers

**DOI:** 10.1101/2021.01.28.428690

**Authors:** Shankar Lalitha Sridhar, Jeffrey Dunagin, Kanghyeon Koo, Loren Hough, Franck Vernerey

## Abstract

Cells are known to use reversible binding to active biopolymer networks to allow diffusive transport of particles in an otherwise impenetrable mesh. We here determine the motion of a particle that experiences random forces during binding and unbinding events while being constrained by attached polymers. Using Monte-Carlo simulations and a statistical mechanics model, we find that enhanced diffusion is possible with active polymers. However, this is possible only under optimum conditions that has to do with the relative length of the chains to that of the plate. For example, in systems where the plate is shorter than the chains, diffusion is maximum when many chains have the potential to bind but few remain bound at any one time. Interestingly, if the chains are shorter than the plate, we find that diffusion is maximized when more active chains remain transiently bound. The model provides insight into these findings by elucidating the mechanisms for binding-mediated diffusion in biology and design rules for macromolecular transport in transient synthetic polymers.

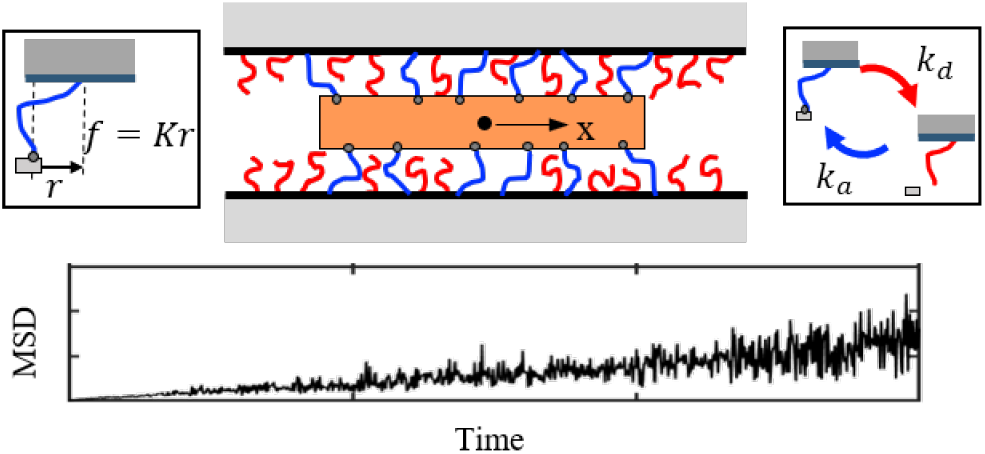

## Introduction

The diffusion of macromolecules in polymeric systems is a key player in self-healing,^1^ biological sorting, recognition^2,3^ and growth.^4^ The physics that drives this transport depends on both the nature of the polymer and the diffusing particles. In covalent networks, transport is usually controlled by the polymer mesh size such that particles with smaller radius of gyration easily diffuse while their larger counterparts are limited to sub-diffusive behaviors.^5^ For physically cross-linked polymers such as alginates, collagen or agarose, the continuous association and dissociation of network junctions not only creates opportunities for diverse mechanical responses^6–8^ but also for enhanced diffusion.^9^ The situation becomes more interesting when diffusing particles have an affinity for the polymer, enabling them to selectively bind and unbind with the surrounding chains and further facilitate diffusion. This affinity-based, rather than size-based mode of diffusion^10^ has been found in a number of biological systems, including the mucus, ^11^ the extracellular matrix^12^ and the nuclear pore complex (NPC), a bio-polymer that acts as a gateway mediating the passage of macromolecules between the cell’s nucleus and its cytoplasm.^9,13–15^ Interestingly, affinity-based motion may also be powered by molecular motors that can convert chemical to mechanical energy in order to speed-up diffusion.^16^ Prominent examples of such motion include intracellular transport of organelles using motor proteins^17^ and the twitching surface motility of bacterial cells that use long polymeric appendages called pili that are powered by motors at their base to actively propel themselves in various environments.^18^

The discovery of these puzzling biological phenomena has motivated experimental research in creating synthetic systems that can perform similar functions in applications such as drug delivery, controlled assembly pathways and tissue engineering.^9,19–21^ Progress in these directions have however been hindered by the lack of a molecular theory for binding-enabled diffusion where particles are driven by unbinding and binding of stretched polymers. Such a theory must help us identify the transition between enhanced diffusion from particle-hopping among neighboring chains and reduced diffusion from the increased “stickiness” of the chains on the particles. It should also clearly define the role of chain length, binding/unbinding kinetics and the size of a particle on this transition. Previous studies on the NPC^22,23^ predict that diffusion is enhanced or hindered based on three time scales namely binding kinetics, elastic relaxation and solvent diffusion.^24,25^ Other models have further explored the sub-diffusive transport of a particle constrained in an energy well by its closest neighbors, modeled as rigid walls.^22,26^ While these studies have enlightened the physics of binding mediated motion, there is still no theory predicting the long-time diffusivity of the particle in terms of molecular parameters.

In this article, we tackle this problem by considering a reduced, one-dimensional model that describes a plate (the particle) diffusing within a population of fluctuating polymer chains via intermittent binding and unbinding events (Fig. 1). In a “passive” system that is under thermodynamic equilibrium, these events act to effectively increase the environment viscosity and consequently decrease the particle’s diffusivity. However, when the chains are pushed out of equilibrium by an internal energy source making them “active”, their fluctuations can be significantly enhanced. This situation can act as a double-edge sword: it can either speed-up the transport or stop it completely by trapping a particle. Our objective is thus to elucidate the role of key physical parameters on plate diffusion, namely (a) the level of activity, (b) the number of binding chains (and corresponding sites), (c) the length of the chains, and (d) the binding/unbinding kinetics. We show here that with optimum combinations of these key parameters, one can enable high particle diffusivity for particles of large size, while trapping smaller particles, which is counter-intuitive to typical diffusive behavior.

**Figure 1:**
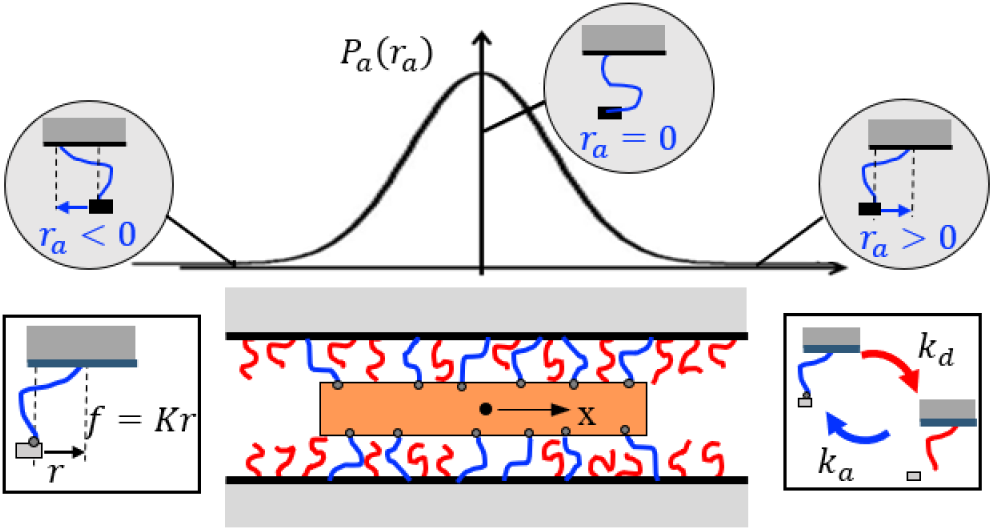
One-dimensional diffusion of plate by reversible binding at rates *k_a_* and *k_d_* to polymer chains grafted on either side. The projection of the chain configuration *r_a_* during binding follows a Gaussian distribution *P_a_*(*r_a_*) and the chains act like linear springs.

### Model setup and Monte Carlo simulations

We present a general model focused on motion in one dimension (*x*) of a plate between two parallel walls grafted with a population of flexible chains that can bind to any location on the plate surface (Fig.1). Each chain undergoes thermal or active fluctuations, and is characterized by *r*, the projection of the chain’s end-to-end vector in the *x* direction. We assume that these fluctuations are Gaussian, and so the quantity r is taken to be normally distributed with mean 0 and variance *σ*^2^ = *N_k_b*^2^/3 where *N_k_* and *b* are the number and length of a Kuhn segments, respectively.^27^ When the end of a chain meets the plate’s surface, it can attach in order to provide a physical connection that acts as a Hookean spring with stretch force *f = Kr*, where *K = k_B_T/σ*^2^ is the equivalent spring stiffness for a given temperature *T*. As a first approximation, we ignore interactions between the chains, excluded volume effects, competition for binding sites, cross-linking of the chains across the plate, and stiffening of the chain as the extension approaches its contour length. These assumptions limit the applicability of our model to situations where the grafting density of chains is sparse enough for inter-chain interactions to be safely ignored. The parameter *N* is introduced to describe the total number of chains in the vicinity of the plate that have the potential to bind. Given a plate length, *L_p_*, and chain grafting density, *ρ*, the total number of attachable chains is *N = ρL_p_*.

Two types of diffusive behavior are considered: the “passive” case where the chains are in thermodynamic equilibrium with the surrounding medium, and the “active” case where they are driven out-of-equilibrium by an internal energy source like motor proteins. We assume that the fluctuations of active chains remain unbiased about the center and reach a nonequilibrium steady state with the medium at long times. Under these assumptions, we then characterize the activity level in the chains by evoking an effective temperature *T_a_*.^28,29^ We note that the original source of the activity in many systems involving motor proteins have a basis in conformation changes and power strokes that produce a conversion of chemical to mechanical energy at a constant temperature.^30^ However, to maintain the simplicity of a freely-jointed polymer model of the chains, we use the effective temperature as a measure to characterize the chain’s active energy. In the passive case, the effective temperature equals that of the medium, *T_a_ = T*, while an active case corresponds to *T_a_ > T*.

Based on this view, we construct a kinetic Monte Carlo scheme of bond dynamics that follows a Poisson process with average rate of attachment, *k_a_*, and detachment, *k_d_*. The lifetime probabilities of the attached state 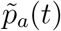 and detached state 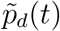 follow an exponentially decaying distribution with a single time scale given by

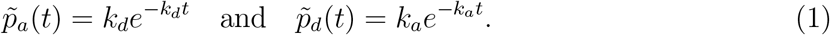

For the passive case, the rates *k_a_* and *k_d_* are given by Eyring’s theory in terms of the temperature *T* and the free energy barrier for the state transition Δ*G*. The frequency of attempts by a chain to attach or detach to the plate is governed by thermal fluctuations as characterized by the Rouse time, 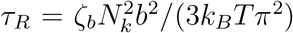, where *ζ_b_* is the local friction coefficient of a chain. Given an attempt frequency *ω* = 1/*τ_R_* œ *k_B_T*, the binding and unbinding rates are, respectively *k_a_ = ω* exp(—Δ*G_a_/k_B_T*) and *k_d_* = *ω* exp(-Δ*G_d_/k_B_T*), where in general Δ*G_a_* = Δ*G_d_*. In the passive case, *ω* is proportional to the medium temperature T as the primary source of energy is from molecular collisions that produces thermal fluctuations of the chain. However, in the active case the attempt frequency is determined by the level of active energy from an extrinsic source. For simplicity, we assume that *ω* for the active case is proportional to the effective temperature of the chain *T_a_*. The potential energy of the active chain is assumed to be encompassed in the effective temperature *T_a_*, which in turn manifests itself in the effective spring constant as *K = k_B_T_a_/σ*^2^. The hydrodynamic effects of the active polymer chain on the medium temperature is out of the scope of this paper and is a subject of interest for future studies.

The probability for a chain to be in the attached state is then given by the equilibrium rate constant *p* = *k_a_*/(*k_a_ + k_d_*). The Monte Carlo step is used to stochastically break current bonds or form new ones based on bond probabilities. Depending on the value of *p*, the steady-state number *N_a_* of chains attached to the plate fluctuates, resulting in a fluctuating force 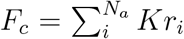 (Fig. 2a). Additionally, the plate is subjected to a fluctuating random force *F_m_*(*t*) due to molecular collisions from the surrounding fluid medium. The time correlation of this force can be obtained from the fluctuation-dissipation theorem as 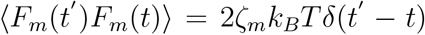, where *ζ_m_* is the plate’s friction coefficient that depends on the medium viscosity *η* and the plate length *L_p_* as *ζ_m_* = 4*πηL_p_*/3.^27^ Assuming the correlations between *F_c_* and *F_m_* to be minimal and neglecting inertial contributions, we balance the forces of drag and thermal fluctuations to calculate the effective plate velocity *v_p_* and the corresponding position *x*(*t*) by integrating in time. We note that this is not true in general, particularly when the hydrodynamic effects on the plate due to the active chains are significant.

**Figure 2:**
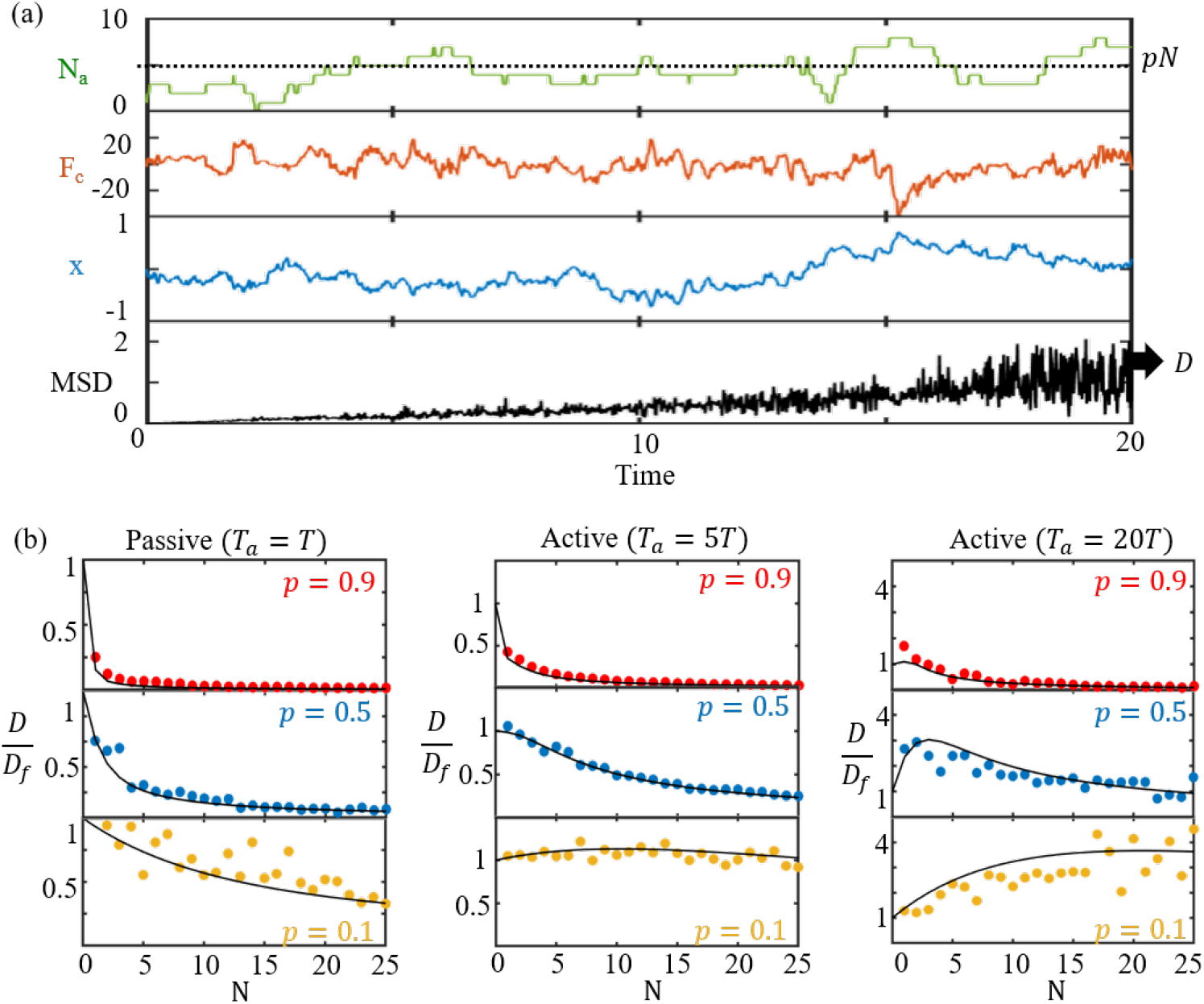
(a) A typical example of binding and unbinding shown as a plot in time of the number of attached bonds *N_a_*, the force on the plate *F_c_* and the position *x* of the center of the plate (*N* = 10,*k_a_/k_d_* = 1,*σ* = 0.17). (b) Plot of diffusion coefficient normalized to the free diffusivity *D_f_* from simulations (circles) and the analytical model (curves) w.r.t the total number of chains *N*.

For each simulation, we calculate the mean squared displacement using a temporal moving average 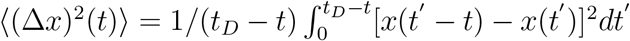, where *t_D_* >> *t* is the total duration of the experiment.^31^ The effective long time Fickian diffusion constant *D* = 〈(Δ*x*)^2^〉/2*t* is then obtained by averaging several simulations each conducted for various combinations of *N, k_a_, k_d_* and *σ*^2^. The effective diffusion constant is normalized as *D*\* = *D/D_f_*, where *D_f_ = k_B_T/ζ_m_* is the free diffusivity due to molecular collisions from the surrounding fluid. Fig. 2b depicts the results of dimensionless diffusivity D* for the passive (*T_a_ = T*) and active cases (*T_a_* = 5*T* and *T_a_* = 20*T*) as a function of the equilibrium constant *p* and the number of available chains *N*. The values of *T_a_* considered here represent the magnitude of free energy (≈ 10 — 30 *k_B_T*) arising due hydrolysis of ATP, a process common in biological systems like bacteria or eukaryotic cells.^32–34^ The contour length of the chain *l_c_ = N_k_b* and the plate length *L_p_* also play a role in diffusivity of the plate by mediating the friction from chains and the medium, respectively. It can be shown that *l_c_/L_p_* ∝ *τ_R_/τ_b_*, the ratio of Rouse time to bound time *τ_b_ = ζ_m_/K* of a Brownian particle in a harmonic well (see supplementary information). We choose for illustration a value of *τ_R_/τ_b_* = 0.8 in Fig. 2 which corresponds to roughly *l_c_ ≈ 2L_p_*, that has a relevant order of magnitude in biological systems like the nuclear pore complex, cytoskeleton, or bacterial motility.^17,18,35^ The role of this parameter is discussed in more detail later.

In the passive case, the effective diffusion coefficient monotonically decreases with both the number of attachable chains or binding sites *N* and the equilibrium constant *p*. We find that the maximum diffusivity in this case is the intrinsic free diffusivity *D_f_* that occurs when hardly any chains are bound – i.e., when either *p* → 0 or *N* → 0. In the active case, however, there is a maximum in *D* for intermediate values of *p* or *N*. At *T_a_* = 5*T*, a transition is observed where the magnitude of diffusivity remains unaffected even when chains are bound. Above this transition temperature, diffusivity is enhanced significantly more than free diffusivity *D_f_* showing a clear peak for *T_a_* = 20*T* at non-zero values of *p*. When *p* → 1, the chains become too sticky and remain permanently attached to the plate, resulting in a zero long term diffusivity for all cases.

### Statistical mechanics model of diffusion

To better explain and predict these trends, we now build a one-dimensional molecular theory (Fig. 3). The plate experiences two types of fluctuating forces arising from (a) the surrounding medium, and (b) the chain attachments and detachments. Because these forces are independent, we begin with the isolated response of the plate solely due to chain fluctuations. We represent the location *x* of the plate at time *t* by the probability density *P(x,t),* for a given number *N_a_* of attached chains. If *w*(*δ, N_a_*) is the stochastic rate of moving this plate by a displacement *δ* due to intermittent binding/unbinding events, the evolution of *P* follows the master equation:

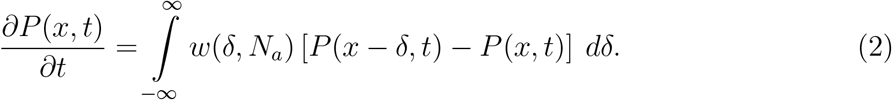

**Figure 3:**
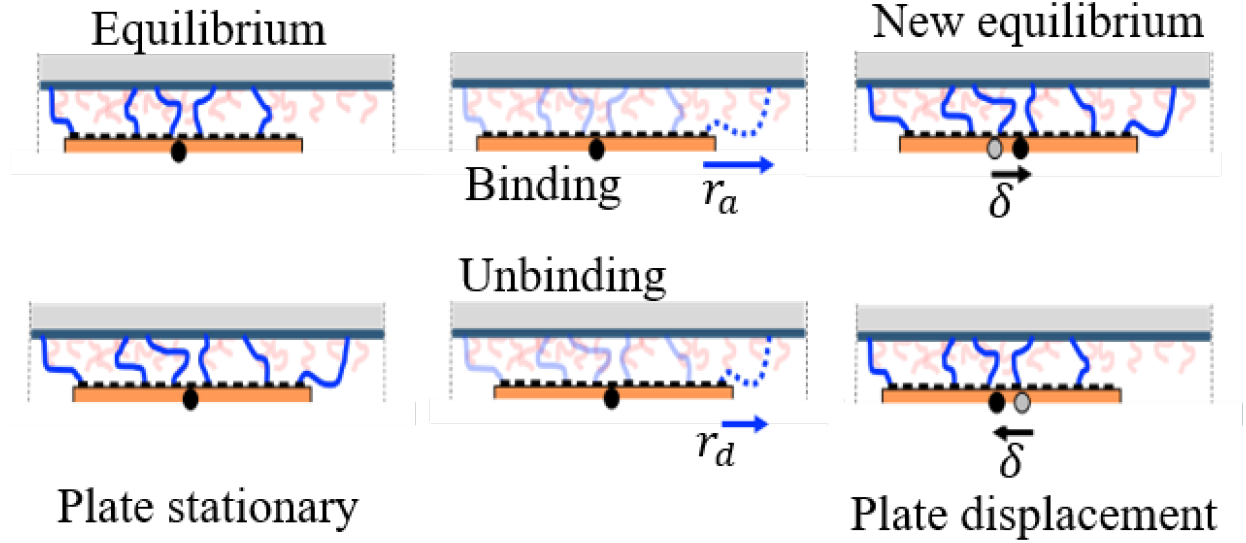
Illustration of the mechanism of plate displacement *δ* and mechanical equilibrium during a new binding or unbinding event.

In other words, the function *P*(*x,t*) decreases when an event causes the plate to leave position *x*, and increases when an event moves the plate from *x – δ* to *x*. The stochastic rate function *w*(*δ, N_a_*) represents the cumulative effect of chain attachments and detachments taking the form

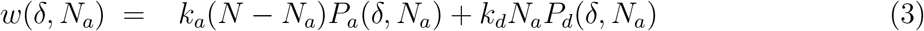

where *P_a_*(*δ, N_a_*) and *P_d_*(*δ,N_a_*) denote the probability density functions (pdf) of the chain configuration during an attachment and detachment event respectively that produces a plate displacement *δ.* As the chains are assumed to follow Gaussian behavior, we assume pdfs for *delta* of the form,

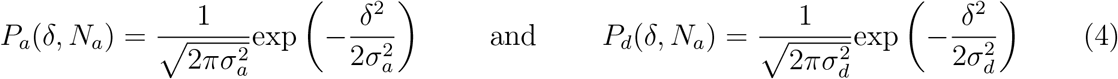

where *σ_a_* and *σ_d_* are the corresponding standard deviations for attachment and detachment events that we now show depend on the number of attached chains *N_a_*. To complete our model, we assume that the pdfs are determined by enforcing force equilibrium on the plate before and after each event. As the plate is in equilibrium with *N_a_* bound chains, an attachment event introduces a new force *F* = *Kr_a_*, where *r_a_* is the lateral projection of the end-to-end vector of the newly attached chain. This immediately triggers a plate displacement *δ = r_a_*/(*N_a_* + 1) to re-establish equilibrium as shown in Fig. 3. Using a similar argument, we find that the plate displacement resulting from the detachment of a chain at configuration *r_d_* is given by *δ = r_d_*/(*N_a_* — 1). The probability distributions of *r_a_* and *r_d_* are Gaussian with a zero-mean and variance 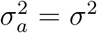 (given simply by the chain statistics) and 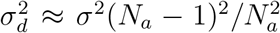 respectively. The latter is obtained by correcting for change in the configurations of attached chains due to the displacement *δ* upon attachment.

Combining 2 and 3, we can characterize the diffusive nature of the plate for an initial state of *N_a_* attached chains. As shown in supplementary information, the plate’s mean displacement 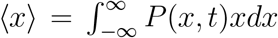 vanishes. Using the mean square displacement, 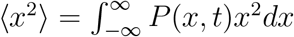, one can derive its corresponding evolution equation from (2). This allows us to determine the plate diffusivity solely due to chain fluctuations (for a given number of attached chains *N_a_*) as 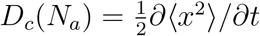, leading to the closed-form solution:

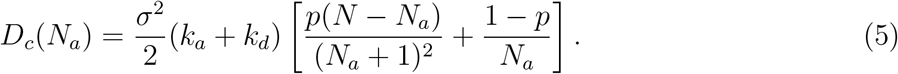

This result shows that the mean-squared displacement is a linear function of time with a constant long-term diffusion coefficient for a given set of input parameters, which implies that the fluctuation-dissipation theorem (FDT) should be satisfied.^22^ Therefore, when considering long timescales, the time correlation of the fluctuating plate force for a given *Na* can be approximated as 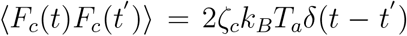, where *ζ_c_ = k_B_T_a_/D_c_*(*N_a_*) is the effective friction coefficient due to chain binding events. Though the chains are out of equilibrium with their surroundings, the use of FDT with an effective temperature *Ta* can be justified under the conditions that the system is at steady state at times much longer than 1/(*k_a_* + *k_d_*).^16,28,29^

The Langevin equation for the plate consists of two types of drag forces and their corresponding force fluctuations which can be written as (*ζ_m_* + *ζ_c_*)*v_p_*(*t*) = *F_c_*(*t*) + *F_m_*(*t*). The effective diffusivity can then be obtained from the velocity correlation of the plate as 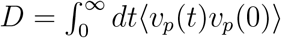. Using the Langevin equation and the FDT with corresponding temperatures, *T* and *T_a_*, we obtain the effective diffusivity as (see supplementary information)

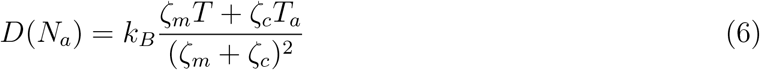

where the effect of correlations between the random forces *F_m_* and *F_c_* on the plate diffusivity are assumed to be negligible. For large values of *N*, 〈*N_a_*〉 = *pN* is a close approximation for the average number of chains and the formula of diffusivity in Eq. 6 is a good estimate. For small values of *N*, however, large fluctuations in the value of *N_a_* become important and is characterized by the binomial distribution 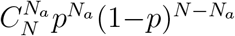 where 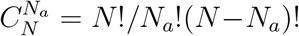. In the general case, the average plate diffusivity can then be calculated as the weighted sum of *D*(*N_a_*) with *N_a_* ∈ (0, *N*), i.e.:

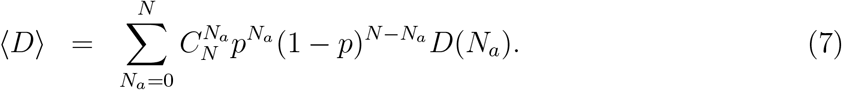

## Results and Discussion

The predictions of Eq. 7 are compared with the simulation results in (Fig. 2b), showing excellent agreement. To gain a general understanding of the model predictions, we turn to the analytical form of Eq. 6. Let us define the following non-dimensional parameters: diffusion constant *D*\* = *D*(*N_a_*)/*D_f_*, friction *ζ*\* = *ζ_c_/ζ_m_*, and temperature *T*\* = *T_a_/T*. The effective diffusion constant from Eq. 6 can now be reorganized into

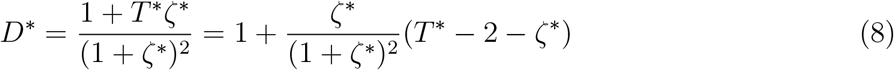

where *ζ*\* can be obtained using the relations *τ_R_* = 1/(*k_a_* + *k_d_*), *K = k_B_T_a_/σ*^2^, and *τ_b_ = ζ_m_/K* as

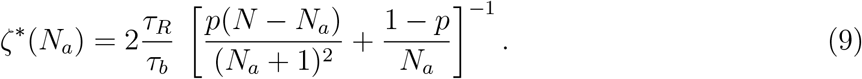

We first recognize that *τ_R_/τ_b_ = ζ_R_/ζ_m_* measures the relative Rouse friction of the chain due to the medium w.r.t medium friction on the plate. This ratio is shown to be proportional to *l_c_/L_p_*, where *l_c_ = N_k_b* and *L_p_* are the lengths of the chain and plate respectively (see supplementary information). The second term in the product of Eq. 8 is interpreted as a correction factor to account for resistance to plate movement by bound chains. When all chains are attached (*p* → 1), the correction factor approaches infinity that results in the effective diffusivity approaching 0. When no chains are bound, (*p* → 0), we find *ζ*\* → 0 and the effective diffusivity *D → D_f_*.

### Threshold temperature for enhanced diffusion

We observe from Eq. 8 that to boost diffusivity (*D*\* > 1), the necessary condition is *T*\* > 2 + *ζ*\*. In other words, enhanced diffusion is possible only if the active temperature *T_a_* is at least twice that of the medium temperature. While the mere presence of activity in the chains (*T_a_* > *T*) will increase the effective diffusivity of the plate compared to a passive system *(T_a_ = T*), it cannot surpass the free diffusivity *D_f_* unless it satisfies the necessary condition described above. Secondly, because of the fluctuation-dissipation theorem, the energy from kicks to the plate due to the medium (*T*) and active chains *(T_a_)* are weighted by their corresponding friction constants *ζ_m_* and *ζ_c_*, respectively (Eq. 6). This implies that diffusivity can be maximized when the friction *ζ_c_* is optimum. An extremely low *ζ*\* can reduce the overall dissipation but it also reduces the weight on the active kicks. If *ζ*\* is too high, the weight on active kicks may be large but the overall resistance can prove too high.

To illustrate the above aspects of diffusion, we identify four dimensionless parameters, *T_a_/T, τ_R_/τ_b_, N*, and *p*, that determine the effective diffusivity of the plate. Each of these parameters represent one physical property of the system: active energy, relative chain to plate lengths (*l_c_/L_p_*), number of attachable chains or binding sites, and relative binding/unbinding rates. We explore the roles of each parameter by presenting two sets of model predictions of diffusivity as a function of *N* and *p* with fixed (i) *τ_R_/τ_b_*, and (ii) *T_a_/T*.

### Non-monotonicity due to active temperature

In the first example, the ratio *τ_R_/τ_b_* is fixed at a value of 0.8 which is so chosen to represent the relevant range of length scales *l_c_* = 1 — 2 *L_p_* in biological systems like the nuclear pore complex.^35,36^ Fig. 4 depicts the plot of effective diffusivity in terms of *N* and *p* for three different chain temperatures with (a) *T_a_ = T*, (b) *T_a_* = 5*T*, and (c) *T_a_* = 20*T*. In the passive case with *T_a_ = T*, the maximum diffusivity is the free diffusivity of the plate *D_f_* and occurs when there are no bound chains. However, in the active cases of *T_a_* = 5*T* and *T_a_* = 20*T*, we find a transition into non-monotonicity seen before from simulations in Fig. 2b. While in the case of *T_a_* = 5*T*, the increase in diffusivity over *D_f_* is only marginal, it is more than three times in the case of *T_a_* = 20*T*, showing the critical role of *T_a_* in enhancing diffusivity. This indicates that *T_a_* = 5*T* is just over the critical temperature 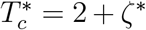 needed for surpassing free diffusivity. The plots in Fig. 4c further show that as *N* increases, the maximum diffusivity occurs at increasingly lower values of *p*, consistent with our simulation results. Moreover, we find that the peaks approach a global maximum *D_M_* asymptotically when *N* → ∞. Thus, the highest diffusivity in this regime occurs when there are infinitely many available chains for binding (*N* → ∞), but have a weak affinity, i.e., unbinding is much faster than binding, such that a few chains are attached to the plate at any given time. This particular combination yields the highest frequency of kicks to the plate (see Eq. 3) which when combined with few chains resisting plate motion gives us the maximum limit in diffusivity. Although the assumption of negligible correlations among different chains in this model are susceptible to breakdown in these limits, this finding provides a valuable insight and a qualitative trend of the maximum diffusivity regime.

**Figure 4:**
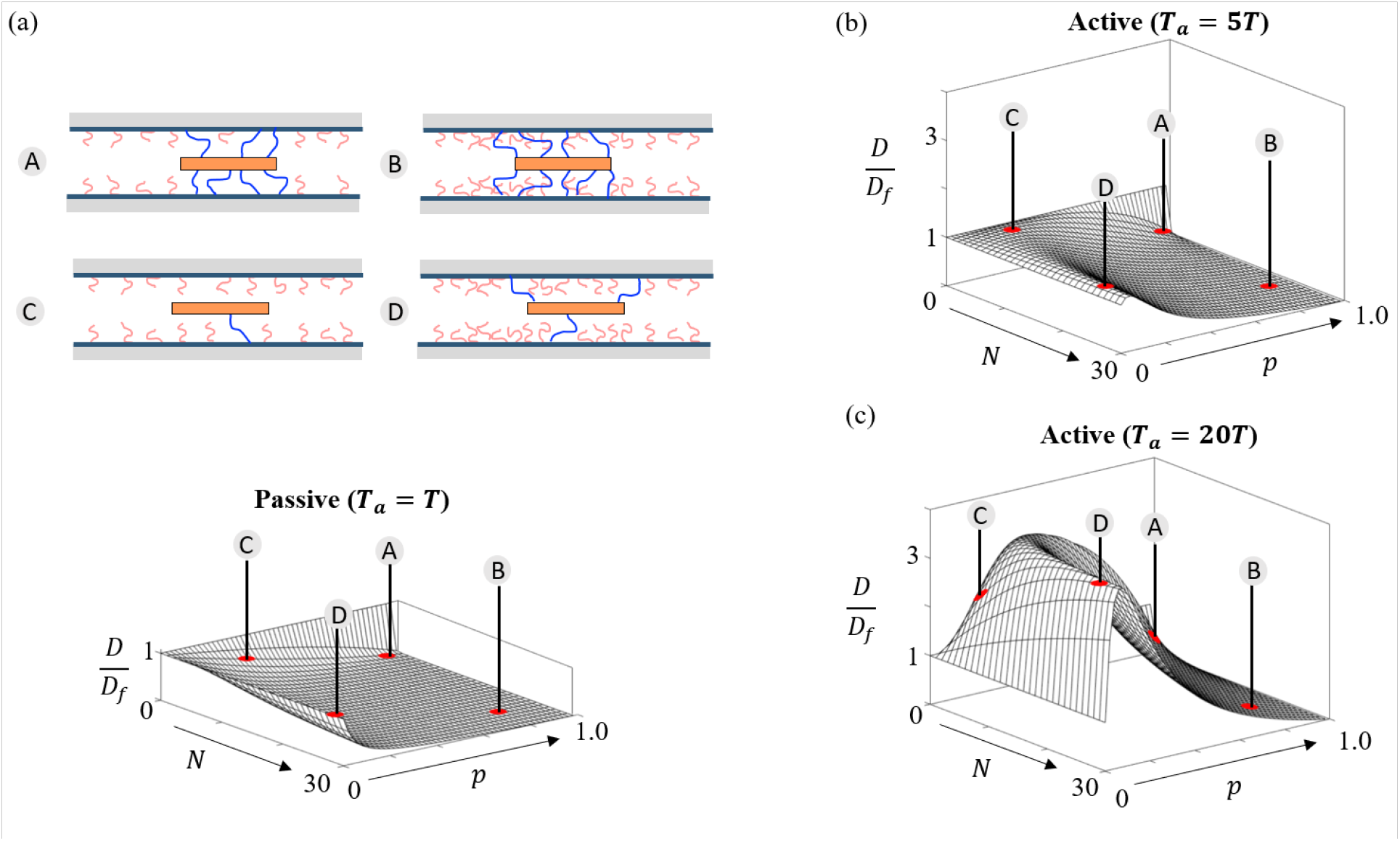
Plots of the normalized diffusion coefficient *D/D_f_* plotted against the total number of chains *N* and rate constant *p = k_a_*/(*k_a_* + *k_d_*). Four different combinations of *p* and *N* are schematically illustrated and categorized A, B, C and D. (a) Passive case with *T_a_ = T*, and (b) Active case with a higher effective chain temperature *T_a_* = 5*T*, and (c) Active case with chain temperature *T_a_* = 20*T*. The ratio of the timescales *τ_R_/τ_b_* = 0.8.

At *N* = 0 and *p* → 0, the transport process reverts back to that of an unbound particle with an intrinsic diffusivity *D_f_*, while for *p* → 1 or *N* → ∞ for a finite *p*, the particle is trapped and its diffusivity vanishes. This type of non-monotonic dependence of diffusivity has been observed with previous models of transport in the nuclear pore complex and mucous membranes, although the model construction and underlying assumptions are different.^22,25,35^ Interestingly, the non-monotonicity in diffusion is also seen in synthetic systems of multi-valent ligand molecules that consists of polyethylene glycol chains (“legs”) that walk or hop by binding to receptor surfaces.^37^

### Effect of chain/plate length on maximum diffusivity

For the second example, we now explore the role of the ratio of chain to plate length, *l_c_/L_p_* ∝ *τ_R_/τ_b_*. With the active temperature fixed at *T_a_* = 20*T*, two new scenarios are considered as shown in Fig. 5a with *τ_R_/τ_b_* = 0.01 and 5b with *τ_R_/τ_b_* = 10. We find an interesting trend where the maximum diffusivity in Fig. 5a, which surpasses that of Fig. 4c with *τ_R_/τ_b_* = 0.8, is now located at a higher *p*, i.e., a larger fraction of bound chains. In other words, when the plate is much longer than the chains (low *τ_R_/τ_b_*), the diffusivity is maximum when more chains are bound as illustrated by case *β* in Fig. 5. By contrast, when the chains are much longer than the plate (case *α*), as in Fig. 5b, we see that there is no enhancement of diffusivity in spite of the presence of active chains and the plot resembles that of passive diffusion *(T_a_ = T*). These results are further corroborated by Monte Carlo simulations presented in the supplementary information (Fig. S1). Projecting the curves in Figs. 4c, 5a, and 5b with *T_a_* = 20*T* to a fixed number of available chains *N* = 20, we find that *τ_R_/τ_b_* dictates the the optimum combination of binding rates (*p = k_a_*/(*k_a_* + *k_d_*)) for maximum diffusivity. As shown in Fig. 5c, the peaks of the effective diffusivity transitions from *p* = 0 for *τ_R_/τ_b_* = 10 to progressively higher values of *p* for smaller values of *τ_R_/τ_b_* = 0.8 and 0.01.

**Figure 5:**
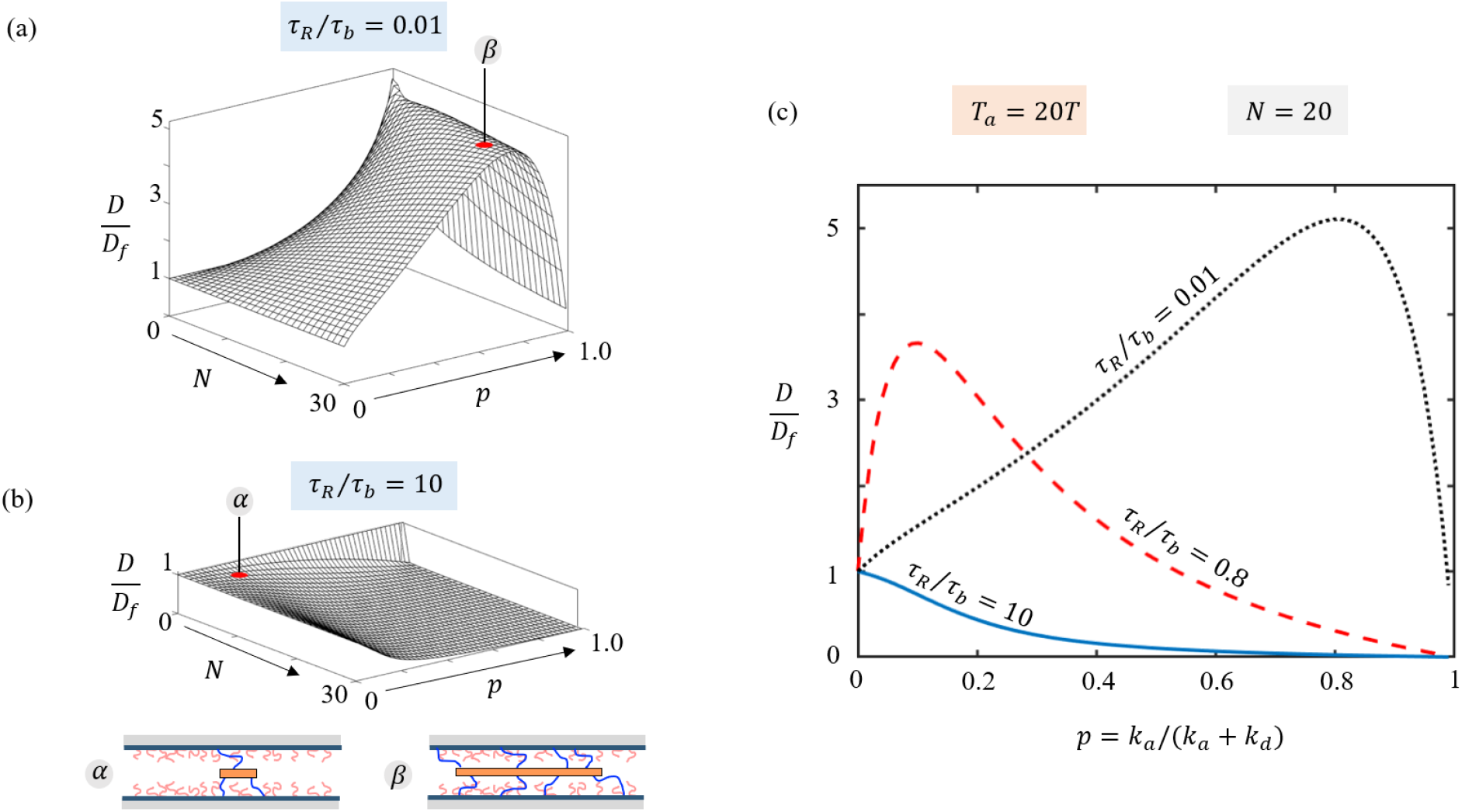
Analysis of the diffusion coefficient for fixed *T_a_* = 20*T* as a function of the ratio of timescales *τ_R_/τ_b_*, which represent the Rouse time of the chains and the bound time of the plate in the fluid medium. Plots of *D/D_f_* for (a) Low (*τ_R_/τ_b_* = 0.01) with peak at point *β*. (b) High *(τ_R_/τ_b_* = 10), presented as functions of total available chains *N* and the fraction of bound chains *p = k_a_*/(*k_a_* + *k_d_*). (c) Cross-sectional slice of 3D plots for *N* = 20 in Figs. 5a, 5b and 4c to show diffusivity as a function of *p*.

To better understand the role of *τ_R_/τ_b_*, we plot the approximate value of maximum diffusivity *D_max_* as a function of *τ_R_/τ_b_* (Fig. 6a). Two key features are notable. First, *D_max_* remains steadily above the free diffusivity *D_f_* for very low values of *τ_R_/τ_b_* ≪ 1 before starting its decline as it approaches the limit *τ_R_/τ_b_* → 1. The magnitude of enhanced diffusivity is larger with higher active temperatures *T_a_*, as expected. Second, we find that depending on the active temperature *T_a_*, there is a critical value of *τ_R_/τ_b_* above which there is no enhancement, i.e., *D_max_/D_f_* = 1. Interestingly, we find that the optimum conditions for *D_max_* in terms of the fraction, *p*, of bound chains is predicted to increase from *p* → 0 at large values of *τ_R_/τ_b_* to *p* → 1 at smaller values (Fig. 6b). For extremely low values of *τ_R_/τ_b_* due to short chains *(l_c_ ≪ L_p_*), the model assumptions like Gaussian chain or one-dimensional behavior may no longer hold.

**Figure 6:**
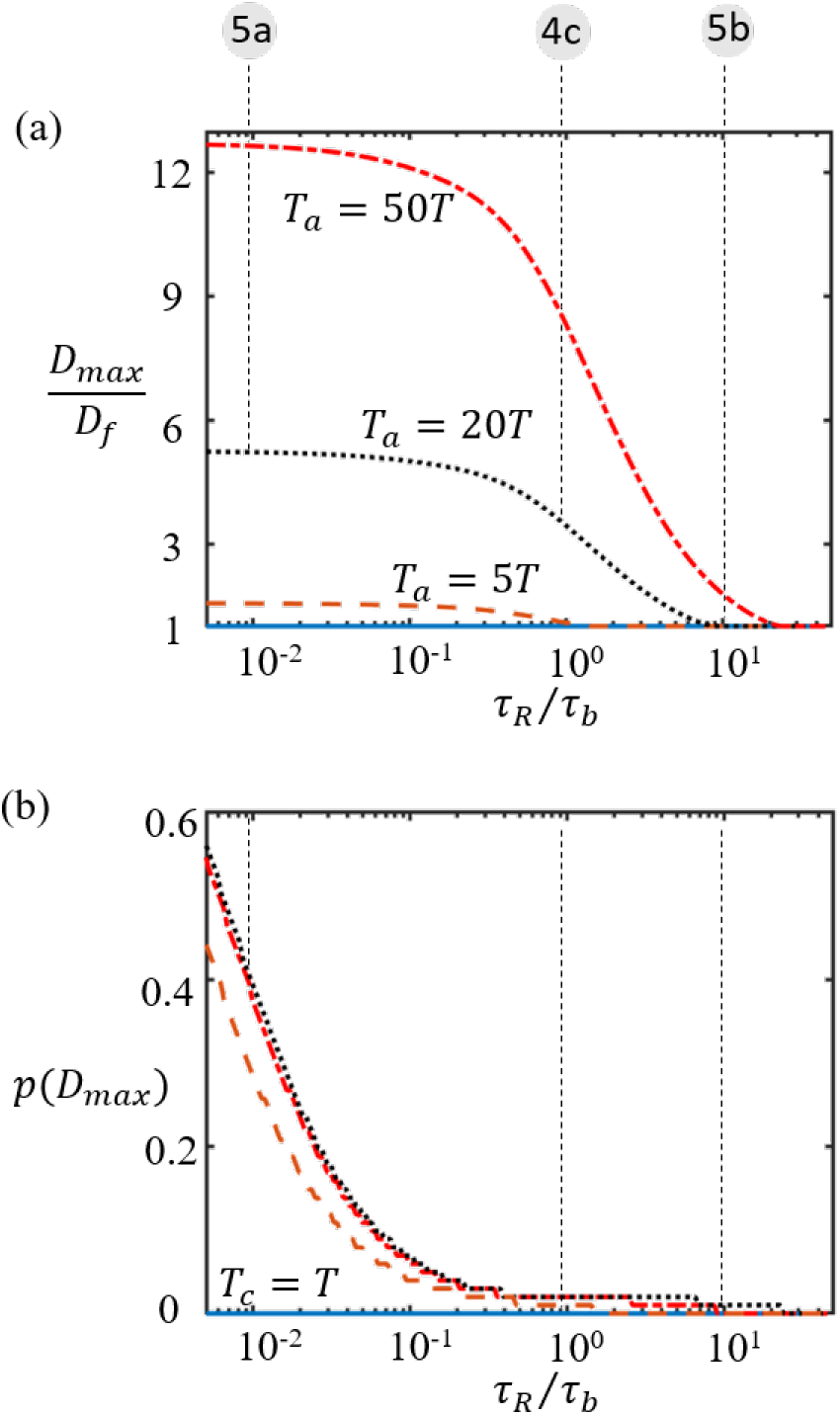
(a) The approximate asymptote of the maximum diffusivity *D_max_* is plotted as a function of *τ_R_/τ_b_* for active temperatures *T_a_ = T, T_a_* = 5*T*, *T_a_* = 20*T*, and *T_a_* = 50*T*. (b) The fraction of bound chains *p* at *D_max_* for all 4 active temperatures. The three vertical lines on the plot correspond to the parameters (*T_a_* and *τ_R_/τ_b_*) used in Figs. 4c, 5a and 5b.

### Physical interpretation of model predictions

The modeling framework presented in this work provides new insights and a better understanding of the mechanisms behind binding-mediated active transport. When *τ_R_/τ_b_* is low due to either small chains or a long plate (*l_c_ < L_p_*), the relative friction *ζ*\* is considerably reduced. However, this also lowers the intensity of active kicks on the plate. For instance, if the plate is extremely long, the friction from the medium becomes so high that the displacement due to a kick from the medium is too small compared to that of moving towards a new equilibrium state following binding events. Particularly, having a slower unbinding rate provides more time for the plate to move to its new equilibrium position following a binding event before the newly attached chain unbinds. The model, therefore, predicts that diffusion is highest when most chains are bound, e.g., high *p*. This explains the shift to higher values of *p* for maximum diffusivity in Fig. 6b as *τ_R_/τ_b_* is reduced. This is likely to be the case for transport of large cell organelles through the cytoplasm of cells, like the centering of ooctye nucleus in the cytoplasm powered by motor proteins.^38^

As *τ_R_/τ_b_* increases, i.e., when each chain becomes more comparable or longer than the plate *(l_c_ ≥ L_p_*), the effective friction on the plate can be reduced by having very few chains bound at any given time (low *p*). For a given *T_a_*, going beyond a threshold value of *τ_R_/τ_b_* will make the chain friction so large that the highest diffusivity occurs only when no chains are bound, which is simply *D_f_*. This suggests that if the plate length *L_p_* is too small compared to the chain length *l_c_*, the free diffusivity *D_f_* is already high enough that the active energy of the chains needs to be above a critical value to produce enhanced diffusion. Alternately, of the chains are much longer than the plate, the time between binding events is too long for the plate to reach a new equilibrium position before the next event. In other words, the fluctuations from the chains are too spread apart, and subside between binding events, to produce enhanced diffusion. We note that this model predicts that the minimum active temperature *T_c_* needed to transition to enhanced diffusivity increases linearly with *τ_R_/τ_b_* (see supplementary information, Fig. S2). Therefore, to produce enhanced diffusion, it is not only important to have active chains but the right level of activity for given geometric (*l_c_/L_p_*) constraints.

## Summary and Conclusions

In summary, our analysis suggests that in contrast to passive diffusion there are different regimes in affinity-based active diffusion that can be finely tuned with a few physical parameters. For example, if binding affinity is strong (*k_a_* > *k_d_*), highly multivalent or large molecules (high *N*) are significantly slowed down (case B in Fig. 4) and trapped compared to molecules with lower valencies (case A). By contrast, for weak affinities (*k_a_ < k_d_*), diffusion of highly multivalent or large molecules (case D) can be much higher than those with lower valencies or size (case C). In fact, our findings indicate that high multivalency but low affinity of each binding site is the best case scenario for rapid transport, a situation found in the nuclear pore complex.^39^ The above findings hold true as long as the plate and chain lengths and their corresponding friction constants are comparable *(l_c_* ≈ 1 — 2 *L_p_*). Extending the model analysis to other combinations of the ratio *l_c_/L_p_* œ *τ_R_/τ_b_*, we find that plates that are considerably larger than the chains can move faster when bound to many active chains. This regime produces non-classical diffusion behavior where larger molecules can diffuse faster than smaller ones when the chains are highly active. We also find that the model and simulations predict the non-intuitive phenomenon that the level of activity in the chains need to be more than twice the thermal energy in the medium (*T_a_* > (2 + *ζ*\*)*T*) to surpass free diffusivity. Since the friction from chains is much higher in the case of large values *l_c_/L_p_*, we find the existence of a critical active temperature *T_c_* below which enhanced diffusivity is not possible.

These findings can provide useful insights into the motility of bacteria that use motorized polymers (pili) and the active mechanisms underlying intracellular organelle transport using motor proteins.^17,18^ It could also prove meaningful for better understanding and tuning of highly selective transport of macromolecules through dynamic polymer networks such as the NPC, mucous membranes and extracellular matrix that depend on both size and chemical affinity. The extension of this approach to two and three-dimensional networks will require the inclusion of network topology, the effect of mesh size (relative to particle size) as well as particle geometry. More work in the future will be required to incorporate complex relationships between bond kinetics and chain forces due to structural changes in the polymer (e.g., protein unfolding), bond type (e.g., slip, catch bonds) or motor response, can produce anomalous behavior like subdiffusion or superdiffusion seen in biological systems.^40–42^

## Acknowledgement

FJV gratefully acknowledges the support of the National Science Foundation under Award No. 1761918. LEH acknowledges support of NIH R35 GM119755, NSF 1943488 and NSF DMR-1551095. The content is solely the responsibility of the authors and does not necessarily represent the official views of the National Science Foundation.

## Supporting Information Available

The following files are available free of charge.

- *Supplementary_active_dif f 1.pdf*: This document contains the mathematical derivations of the effective diffusivity from reversible binding and the reduced forms under a high temperature limits.

## References

(1) Stukalin, E. B.; Cai, L.-H.; Kumar, N. A.; Leibler, L.; Rubinstein, M. Self-healing of unentangled polymer networks with reversible bonds. Macromolecules 2013, 46, 7525–7541.

(2) McCammon, J. A. Theory of biomolecular recognition. Current opinion in structural biology 1998, 8, 245–249.

(3) Chou, C.-F.; Bakajin, O.; Turner, S. W.; Duke, T. A.; Chan, S. S.; Cox, E. C.; Craighead, H. G.; Austin, R. H. Sorting by diffusion: An asymmetric obstacle course for continuous molecular separation. Proceedings of the National Academy of Sciences 1999, 96, 13762–13765.

(4) Akalp, U.; J. Bryant, S.; J. Vernerey, F. Tuning tissue growth with scaffold degradation in enzyme-sensitive hydrogels: a mathematical model. Soft Matter 2016, 12, 7505–7520.

(5) Lustig, S. R.; Peppas, N. A. Solute diffusion in swollen membranes. IX. Scaling laws for solute diffusion in gels. Journal of Applied Polymer Science 1988, 36, 735–747.

(6) Vernerey, F. J.; Long, R.; Brighenti, R. A statistically-based continuum theory for polymers with transient networks. Journal of the Mechanics and Physics of Solids 2017, 107, 1–20.

(7) Sridhar, S. L.; Vernerey, F. J. Mechanics of transiently cross-linked nematic networks. Journal of the Mechanics and Physics of Solids 2020, 104021.

(8) Shen, T.; Vernerey, F. J. Rate-dependent fracture in transient networks. Journal of the Mechanics and Physics of Solids 2020, 104028.

(9) Goodrich, C. P.; Brenner, M. P.; Ribbeck, K. Enhanced diffusion by binding to the crosslinks of a polymer gel. Nature Communications 2018, 9, 4348.

(10) Witten, T. A.; Cohen, M. H. Crosslinking in shear-thickening ionomers. Macromolecules 1985, 18, 1915–1918.

(11) Thornton, D. J.; Sheehan, J. K. From Mucins to Mucus. Proceedings of the American Thoracic Society 2004, 1, 54–61.

(12) Zámecník, J.; Vargová, L.; Homola, A.; Kodet, R.; Syková, E. Extracellular matrix glycoproteins and diffusion barriers in human astrocytic tumours. Neuropathology and Applied Neurobiology 2004, 30, 338–350.

(13) Osmanović, D.; Fassati, A.; Ford, I. J.; Hoogenboom, B. W. Physical modelling of the nuclear pore complex. Soft Matter 2013, 9, 10442–10451.

(14) Mincer, J. S.; Simon, S. M. Simulations of nuclear pore transport yield mechanistic insights and quantitative predictions. Proceedings of the National Academy of Sciences of the United States of America 2011, 108, E351–358.

(15) Miao, L.; Schulten, K. Probing a Structural Model of the Nuclear Pore Complex Channel through Molecular Dynamics. Biophysical Journal 2010, 98, 1658–1667.

(16) Loi, D.; Mossa, S.; Cugliandolo, L. F. Non-conservative forces and effective temperatures in active polymers. Soft Matter 2011, 7, 10193–10209.

(17) Soppina, V.; Rai, A. K.; Ramaiya, A. J.; Barak, P.; Mallik, R. Tug-of-war between dissimilar teams of microtubule motors regulates transport and fission of endosomes. Proceedings of the National Academy of Sciences 2009, 106, 19381–19386.

(18) Marathe, R.; Meel, C.; Schmidt, N. C.; Dewenter, L.; Kurre, R.; Greune, L.; Schmidt, M. A.; Müller, M. J.; Lipowsky, R.; Maier, B., et al. Bacterial twitching motility is coordinated by a two-dimensional tug-of-war with directional memory. Nature communications 2014, 5, 1–10.

(19) Lieleg, O.; Ribbeck, K. Biological Hydrogels as Selective Diffusion Barriers. Trends in cell biology 2011, 21, 543–551.

(20) Bryant, S. J.; Vernerey, F. J. Programmable Hydrogels for Cell Encapsulation and Neo-Tissue Growth to Enable Personalized Tissue Engineering. Advanced Healthcare Materials 2017, 7, 1700605.

(21) Sridhar, S. L.; C. Schneider, M.; Chu, S.; Roucy, G. d.; J. Bryant, S.; J. Vernerey, F. Heterogeneity is key to hydrogel-based cartilage tissue regeneration. Soft Matter 2017, 13, 4841–4855.

(22) Bickel, T.; Bruinsma, R. The nuclear pore complex mystery and anomalous diffusion in reversible gels. Biophysical journal 2002, 83, 3079–3087.

(23) Hough, L. E.; Dutta, K.; Sparks, S.; Temel, D. B.; Kamal, A.; Tetenbaum-Novatt, J.; Rout, M. P.; Cowburn, D. The molecular mechanism of nuclear transport revealed by atomic-scale measurements. Elife 2015, 4, e10027.

(24) Fogelson, B.; Keener, J. P. Enhanced Nucleocytoplasmic Transport due to Competition for Elastic Binding Sites. Biophysical Journal 2018, 115, 108–116.

(25) Fogelson, B.; Keener, J. Transport Facilitated by Rapid Binding to Elastic Tethers. SIAM Journal on Applied Mathematics 2019, 1405–1422.

(26) Ribbeck, K.; Görlich, D. Kinetic analysis of translocation through nuclear pore complexes. The EMBO Journal 2001, 20, 1320–1330.

(27) Doi, M.; Edwards, S. F.; Edwards, S. F. The theory of polymer dynamics; oxford university press, 1988; Vol. 73.

(28) Dieterich, E.; Camunas-Soler, J.; Ribezzi-Crivellari, M.; Seifert, U.; Ritort, F. Singlemolecule measurement of the effective temperature in non-equilibrium steady states. Nature Physics 2015, 11, 971–977.

(29) Cugliandolo, L. F.; Kurchan, J.; Peliti, L. Energy flow, partial equilibration, and effective temperatures in systems with slow dynamics. Physical Review E 1997, 55, 3898.

(30) Duke, T. Physics of bio-molecules and cells. Physique des biomolécules et des cellules; Springer, 2002; pp 95–143.

(31) Lubelski, A.; Sokolov, I. M.; Klafter, J. Nonergodicity Mimics Inhomogeneity in Single Particle Tracking. Physical Review Letters 2008, 100, 250602.

(32) Tran, Q. H.; Unden, G. Changes in the proton potential and the cellular energetics of Escherichia coli during growth by aerobic and anaerobic respiration or by fermentation. European journal of biochemistry 1998, 251, 538–543.

(33) Wackerhage, H.; Hoffmann, U.; Essfeld, D.; Leyk, D.; Mueller, K.; Zange, J. Recovery of free ADP, Pi, and free energy of ATP hydrolysis in human skeletal muscle. Journal of applied physiology 1998, 85, 2140–2145.

(34) Philips, R. M. R. » How much energy is released in ATP hydrolysis? http://book.bionumbers.org/how-much-energy-is-released-in-atp-hydrolysis/.

(35) Maguire, L.; Stefferson, M.; Betterton, M. D.; Hough, L. E. Design principles of selective transport through biopolymer barriers. Physical Review E 2019, 100, 042414.

(36) Maguire, L.; Betterton, M. D.; Hough, L. E. Bound-state diffusion due to binding to flexible polymers in a selective biofilter. Biophysical Journal 2020, 118, 376–385.

(37) Perl, A.; Gomez-Casado, A.; Thompson, D.; Dam, H. H.; Jonkheijm, P.; Reinhoudt, D. N.; Huskens, J. Gradient-driven motion of multivalent ligand molecules along a surface functionalized with multiple receptors. Nature chemistry 2011, 3, 317–322.

(38) Almonacid, M.; Ahmed, W. W.; Bussonnier, M.; Mailly, P.; Betz, T.; Voituriez, R.; Gov, N. S.; Verlhac, M.-H. Active diffusion positions the nucleus in mouse oocytes. Nature cell biology 2015, 17, 470–479.

(39) Liu, S. M.; Stewart, M. Structural basis for the high-affinity binding of nucleoporin Nup1p to the Saccharomyces cerevisiae importin-*β* homologue, Kap95p. Journal of molecular biology 20 05, 349, 515–525.

(40) Regner, B. M.; Vučinić, D.; Domnisoru, C.; Bartol, T. M.; Hetzer, M. W.; Tartakovsky, D. M.; Sejnowski, T. J. Anomalous diffusion of single particles in cytoplasm. Biophysical journal 2013, 104, 1652–1660.

(41) Sjollema, J.; van der Mei, H. C.; Hall, C. L.; Peterson, B. W.; de Vries, J.; Song, L.; Jong, E. D. d.; Busscher, H. J.; Swartjes, J. J. T. M. Detachment and successive re-attachment of multiple, reversibly-binding tethers result in irreversible bacterial adhesion to surfaces. Scientific Reports 2017, 7, 4369.

(42) van der Westen, R.; Sjollema, J.; Molenaar, R.; Sharma, P. K.; van der Mei, H. C.; Busscher, H. J. Floating and tether-coupled adhesion of bacteria to hydrophobic and hydrophilic surfaces. Langmuir 2018, 34, 4937–4944.

